# Endosome-ER Interactions Define a Cellular Energy Landscape to Guide Cargo Transport

**DOI:** 10.1101/2023.06.01.543348

**Authors:** Yusheng Shen, Yan Wen, Qirui Zhao, Pingbo Huang, Pik-Yin Lai, Penger Tong

## Abstract

Molecular motor-driven cargo must navigate a complex intracellular environment, which is often crowded, heterogeneous and fluctuating, to fulfill their diverse functions in a cell. To meet these challenges, most cargo display qualitatively similar transport behavior, that is, random switching between states of diffusive “jiggling” movement (off-state) and states of directed “runs” (on-state). The physical picture of this 2-state motion and their regulation in a cell are not well understood. Here, by using single-particle tracking and motion-states dissection, we present a statistical analysis of the 2-state motion of epidermal growth factor receptor (EGFR)-endosomes in living cells. From a thorough analysis of a large volume of EGFR-endosome trajectories, we reveal that their lifetime in both states feature an exponential distribution with its probability density function (PDF)-amplitude varied by more than three decades. We show that their characteristic time, on-state probability, velocity and off-state diffusion coefficient are spatially regulated, and are probably contributed by the endoplasmic reticulum (ER) network via its spatially varying membrane densities and interactions with the cargo. We further propose a 2-state transport model to describe the complex, spatially varying transport dynamics of EGFR-endosomes in a cell. Our findings suggest that the ER network may play an essential role in constructing a cellular-level free-energy landscape Δ*G*(**r**) to spatially guide cargo transport.

---

The endosomal network is an extremely dynamic, yet highly regulated highway system which relies heavily on the molecular-motor-driven transport of cargoes to continually communicate between cellular compartments, acquire nutrients, respond to extracellular signals, and maintain cellular homeostasis [1–3]. Moving along the cytoskeletal tracks consisting of both microtubules (MTs) and actin filaments, on the one hand, cargoes that are carried by motor proteins need undergo directed motion with a characteristic velocity to quickly reach their destination. On the other hand, unloading the cargo to the proper location of function requires a “proofreading” mechanism that cargo transport is guided spatiotemporally by the surrounding environment such that they can “pause”, change directions and search for a correct path to the destination [1, 4, 5].

Although *in vitro* analysis of molecular motors has uncovered many of their intrinsic motion properties such as directionality, run length and velocity [6], cargo transport in living cells is much more complicated than is reflected by the relatively simplified *in vitro* system in which purified motor proteins moving along an individual microtubule. The interior of a living cell, including the tracks, is rather a crowded, heterogeneous and even fluctuating environment [7–14]. First, MTs are decorated with post-translational modifications and non-motile microtubule-associated proteins (MAPs) that could either promote or inhibit the activity of specific motor proteins, and MAPs could act as physical road blocks [15–17]. The cellular distribution of such decorations is highly regulated and some were reported to feature a spatial gradient across the cell [18–20]. Second, the cytoskeletal tracks could be organized into a complex, 3-dimensional network [22, 23]. Bundles could be formed by parallel MTs of either uniform or opposite polarity such as found in the axon and dendrites of mature neurons, respectively [21]. Microtubule intersections can force moving cargoes to slow down, pause, rotate, and change transport directions [24, 25]. Third, oppositely directed molecular motors such as kinesins and dyneins are simultaneously bound to almost all cargoes and transport of the same cargo may involve coordination of motors of varied types and copy numbers [4, 26, 27]. Lastly, cargoes such as endosomes can be tightly tethered to the endoplasmic reticulum (ER) that entangles with the microtubule network and permeates the entire cytoplasmic space [1, 8, 14, 28–30]. A moving endosome will pull ER tubules with them and therefore encounters considerable impedance from the surrounding, heterogeneously distributed ER network [14, 28, 31, 32]. Interestingly, despite such environmental complexity, most cargoes display qualitatively similar transport behaviours, specifically random switching between states of diffusive “jiggling” movement or “pauses” and states of directed “runs” [4, 7, 24, 33– 35]. To analyse and understand such processes within the complex cellular environment is thus fundamental for dissecting the role of individual cellular factors in the regulation of cargo transport.

Due to the complexity, such characterization requires tracking of cargo transport at a high spatiotemporal resolution to allow the identification of distinct states, and meanwhile, large volume of statistics both over space and time to average out random fluctuations in a living cell. However, current analysis of cargo transport, either limited by methodology or by lacking enough statistics, is far from satisfactory. For example, in polarised neuronal cells, movement of individual vesicles in the axon is widely analysed by using kymographs and motion states need to be manually identified [19, 36–38]. This method is time-consuming and normally only tens of trajectories are quantified. Furthermore, spatial resolution is greatly reduced due to the use of segmented lines in generating kymographs, and the detail of the motion dynamics in both states is lost. In non-neuronal cells, single-particle-tracking (SPT) technique and mean-squared displacement (MSD) analysis have been successfully applied to identify and characterize motion states, and have revealed that endosomal motion during the “pause” state is diffusive [7, 33, 39]. However, due to the lack of enough statistics and in-depth analysis of the full spectrum of the motions in these studies especially at the whole-cell scale, a clear physical picture of the motion transitions and their connection to the complex intracellular environment is still missing.

In this study, we focus on the endocytic trafficking of epidermal growth factor receptor (EGFR) in living cells, a well-characterized model for studying the dynamics of endocytic processes and vesicle transport, by using the SPT method. Upon the activation of EGFR by EGF, the EGFR-EGF complex mainly starts their voyage through the endosomal system from early to late endosomes, and are ultimately degraded in lysosomes [40]. The traffic starts from the cell membrane and ends in the perinuclear “cloud” with its routes generally spanning the whole cell. We first choose to investigate the transport dynamics of EGFR-containing endocytic vesicles in BEAS-2B cells, a human lung bronchial epithelial cell line, which typically spread as a thin cell sheet with a large projection area (up to 100 *μ*m × 100 *μ*m) when cultured on a glass coverslip. The thickness of the cell gives us the convenience to largely avoid the out-of-focus problem in most 2D intracellular tracking experiments and meanwhile, allows us to collect a large volume of data to capture the spatiotemporal dynamics of cargo-transport at the whole-cell scale.

From the obtained large volume of individual EGFR-endosome trajectories obtained over a wide range of sampling rates and long durations (total *N >* 469 cells and 70400 trajectories analysed), we identify and extract the “run”/”pause” state segments from each trajectories (on/off state as referred hereafter), and identify a critical parameter, *p*_on_(**r**), of the transport, which features a gradually decrease as the endosome approaches the perinuclear region. Meanwhile, the endosome slow down with a decrease in both on-state velocity and off-state diffusivity. Here, *p*_on_ denotes the on-state probability,

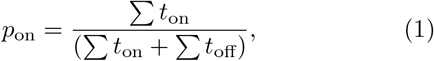

of individual trajectory, where *t*_on_ and *t*_off_ are the duration of the extracted on/off-state segments, respectively. **r** = {*x*}; {*x, y*}; {*x, y, z*} denotes, respectively, the 1*D*, 2*D* and 3*D* spatial coordinates. Interestingly, both *t*_on_ and *t*_off_ are found to be exponentially distributed, suggesting that the transitions between the two states occur randomly in time and can be described by a Poisson process with a constant transition rate *k* = 1*/*⟨*t*⟩ [41]. We propose that the spatially varying transport dynamics are controlled by a cellular-level free free-energy landscape Δ*G*(**r**) which is strongly associated with the spatially varying ER morphology and endosome-ER interactions.

## Results

### Transport of late EGFR-endosomes across the whole cell

To track the motion of EGFR-containing endosomes, EGFRs on the cell membrane are labelled by bright and photostable EGF-conjugated quantum dots (EGF-QDs) at room temperature and the endocytosis of EGFR-EGF-QD containing vesicles (EGFR-endosomes) is monitored as the temperature is elevated to 37 °C. We obtain the EGFR-endosome trajectories from consecutive images of the QDs, and find their position **r**(*t*) (and hence the position of EGFR-endosomes) at time *t* using a homemade single-particle tracking program with a spatial resolution of ∼20 nm. The internalized EGFRs are immunolabeled and are found to be co-localized with EGF-QDs during the endocytosis (Fig. S1), suggesting that the QD-containing vesicles are indeed endocytosed via the EGFR-dependent pathway.

From the trajectories obtained at the whole-cell scale, the EGFR-endosomes are found to be spatially organized into two distinct fractions: a relatively immobile bulk that are trapped in the perinuclear region and a dynamic fraction that occasionally escape to the cell periphery and move bidirectionally along MT tracks (Fig. 1(a and b)). The trajectories of the dynamic fraction displays a general radiating structure around the nucleus that closely resembles the structure of MTs in BEAS-2B cells and are highly heterogeneous with slow-moving endosomes mainly localized at the perinuclear region. Individual endosomes generally exhibit two states of motion, the on-state burst of directed motion and the relatively long, off-state jiggling of diffusive motion (Fig. 1(c)). This is in agreement with previous findings [7, 24, 33–35], however, to our knowledge, here we provide the first picture of such two-state transport dynamics at the whole-cell scale.

**FIG. 1.**
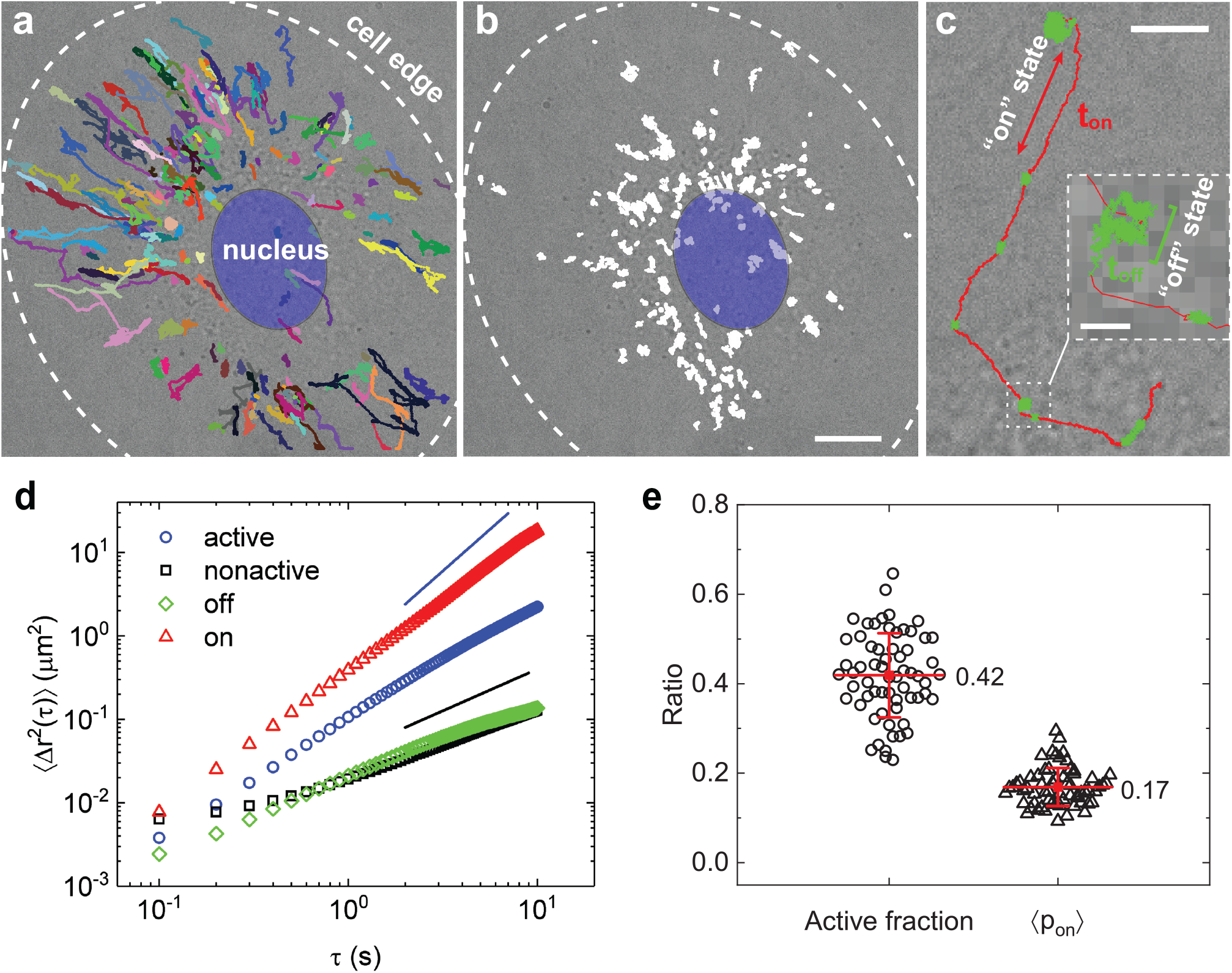
Transport of late EGFR-endosomes across a living cell. (a) 194 representative mobile trajectories (coloured lines) superimposed on a simultaneously taken bright-field image of a living BEAS-2B cell, showing the motion of the EGFR endocytic vesicles inside the cell. The trajectories are taken over 1800 time steps (180-s-long). The white dashed circle indicates the cell periphery and the blue ellipse indicates the cell nucleus. (b) 270 representative nearly-immobile trajectories (white spots) superimposed on a simultaneously taken bright-field image of a living BEAS-2B cell, showing the trapping of the EGFR endocytic vesicles inside the cell. The trajectories are taken over 1800 time steps (180-s-long) and the scale bar is 10 *μ*m. (c) A representative mobile trajectory of a single EGFR endocytic vesicle showing the intermittent switching between the on-state (red) and the off-state (green). Inset shows a magnified view of a portion of the trajectory. The scale bar in (c) is 5 *μ*m and that in the inset is 0.5 *μ*m. (d) Meausred ⟨Δ*r*^2^(*τ*)⟩ as a function of delay time *τ* for the active (blue circles), and non-active (black squares) trajectories, and the on segments (red triangles) and the off segments (green diamonds) of the active trajectories from a single cell. The black and blue solid lines indicate the relationship ⟨Δ*r*^2^(*τ*)⟩ ∼ *τ* and ⟨Δ*r*^2^(*τ*)⟩ ∼ *τ* ^2^ with a slope of unity in the log-log plot, respectively. (e) Cell to cell variation of the ratio of active fraction and mean on-state probability ⟨*p*_on_⟩ for BEAS-2B cells. *N* = 63 cells.

To quantitatively characterize such a highly heterogeneous population of EGFR-endosomes, an automatic algorithm that could characterize the degree of local directionality of movements [42] is applied to identify and extract the on/off (directed/diffusive) state segments from the individual trajectories (see Fig. S2 and Sec. I in Supplementary Materials). From analysing the on-state probability *p*_on_ of individual trajectories, we define the endosomes with *p*_on_ *>* 0, as the dynamic fraction (active,∼ 42%, Figs. 1(a) and (e)), and *p*_on_ = 0, as the relatively static fraction (nonactive, Fig. 1(b)). The dynamic fraction, with a MSD scaling exponent *α* ≃ 1.5, are spatially heterogeneous with *p*_on_ of individual endosomes varying from 0 to ∼0.6 (Fig. 1(d)). Here, *α* is obtained by fitting the MSD data to ⟨Δ*r*^2^(*τ*)⟩ ∼ *τ* ^*α*^. In previous studies, *α* obtained from the local MSD have been used to identify motion states, and trajectory segments with *α >* 1.5 are treated as the on-state [7, 39]. After separating the on- and off-states, the MSD scaling exponent for the off-state segments decreases to *α* ≃ 1, whereas for the on-state segments increases to *α* ≃ 1.7, suggesting our algorithm achieves a sufficient discrimination of motion states under the experimental condition (Fig. 1(d)). The mean on-state probability obtained for the dynamic fraction across different cells is ∼17% (Fig. 1(e)), which is similar to that reported in previous findings [7]. The static fraction are relatively less mobile, and the measured MSD is nearly the same with that of the off-state segments, indicating that this fraction is probably kept in the off state during the observation.

### Characteristic features of the EGFR vesicle trajectories

Because both minus-end directed dynein motors (toward the nucleus) and plus-end directed kinesin motors (toward the cell periphery) are involved in the transport of EGFR-endosomes [9, 40], we perform long-duration tracking up to 1 h to examine how the directionality of transport evolve temporally in individual cells upon the endocytosis of the EGFR-EGF complex (Figs. 2). By comparing the relative distances of the start and end points of each trajectory-segments to the nucleus at given time point, we define the transport direction and quantify the ratio *γ* of vesicles that are directed toward the nucleus to all the vesicles as function of delay time *t*. Two distinct stages of transport are identified: an early stage (*γ >* 0.5) with more vesicles are transported toward the perinuclear region (*t <* 40 min) and a late stage (*γ* ∼ 0.5) with vesicles being transported in both directions at equal probability (*t >* 40 min) (Figs. 2(c)). These two stages probably correspond to the transport of Rab5-dominant early endosomes and Rab7-dominant late endsomes/lysosomes [7, 40], respectively. We next treat the late-stage as a “steady” state for the transport of EGFR endosomes. At this stage, although vesicles are mainly trapped in a broad perinuclear region (Figs. 2(c), inset), they could move bidirectionally across the whole cell, and thus serve as an ideal model for investigating how cellular factors spatially regulate cargo transport.

**FIG. 2.**
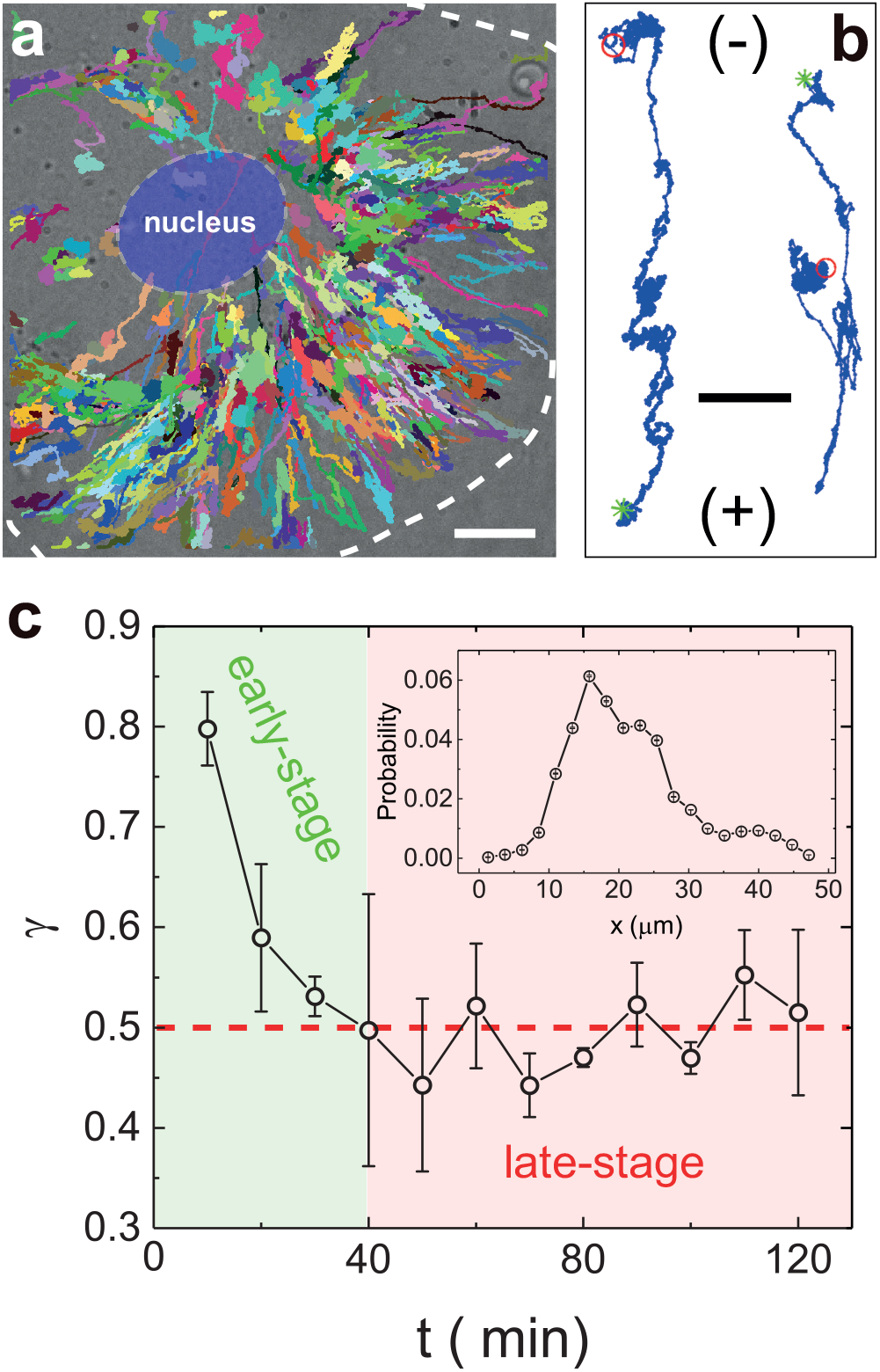
Early-stage and late-stage EGFR-endosomes. (a) 737 representative mobile trajectories (coloured lines) with 4800 time steps (1 h) show the dynamics of EGFR endosomes inside a living BEAS-2B cell. The blue ellipse indicates the nucleus and the white dashed lines indicate the cell edge. (b) A magnified view of two representative trajectories of the early-stage (left) and late-stage (right) EGFR-endosomes. (-) and (+) indicate the minus and plus ends of the MTs, respectively. Green stars and red circles indicate the start and end point of transport, respectively. Scale bars are 10 *μ*m for (a) and 5 *μ*m for (b). (c) Measured ratio *γ* of the EGFR endocytic vesicles moving toward the minus end of the MTs to all the vesicles as a function of delay time *t* upon the endocytosis of the EGF-QDs in a long-duration tracking of EGFR-endosomes. Inset, obtained probability of finding a late-stage endosome as a function of the distance of the endosome to the center of the nucleus. The probability is averaged from the active, late-stage EGFR-endosomes over a 1 h time window from a single cell. The red dashed line indicates the equal probability of minus and plus end directed endosomes observed. The error bar is the standard error of the mean (SEM) across 3 experiments.

Next, we focus on the motion of late-stage EGFR-endosomes and examine how the on-state probability *p*_on_ of individual endosomes evolve spatially. Because *t*_off_ varies in a much wider range than *t*_on_, *p*_on_ is largely determined by *t*_off_. We choose the relative location of off-states, ⟨*ρ*⟩, in the cell as a spatial readout for endosomes with varying *p*_on_. Here, ⟨*ρ*⟩ is the mean of the calculated distance between the nucleus and the off-state sites for a given trajectory. As shown in Figs. 3(b) (black circles), *p*_on_ gradually decrease as the endosome approaches the nucleus, suggesting *p*_on_ and endosome positioning in the off state is spatially regulated by the cell.

**FIG. 3.**
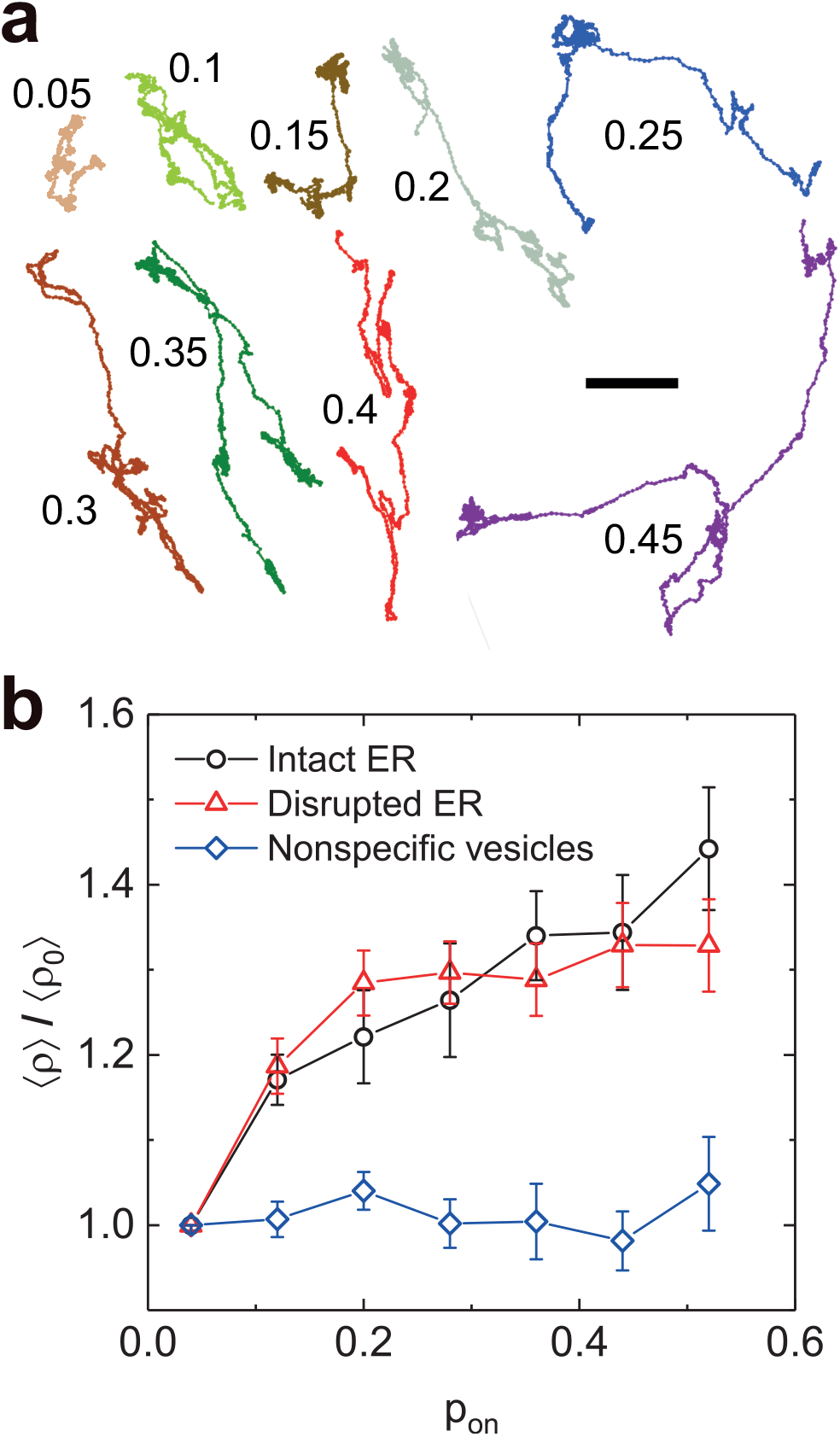
Characterization of the mobile trajectory of EGFR endocytic vesicles. (a) Nine representative mobile trajectories of the EGFR endocytic vesicles for different values of the on-state probability *p*_on_, as indicated by the nearby number. The scale bar is 5 *μ*m. (b) Variations of the measured on-state probability *p*on with varying mean distance ⟨*ρ*⟩ away from the cell nucleus. In the plot, ⟨*ρ*⟩ is normalized by the mean distance ⟨*ρ* _0_⟩ obtained at the smallest value of *p*_on_ = 0.04. The black circles are obtained for EGFR-endosomes with intact ER (*N* = 23 cells), disrupted ER (red triangles, *N* = 46 cells), and Dio-labelled nonspecific vesicles (blue diamonds, *N* = 19 cells). In each cell, ⟨*ρ*⟩ is the mean of the calculated distance between the nucleus and the pausing sites for endosomes of a given *p*_on_ with a bin width of 0.08 and ⟨*ρ* _0_⟩ is the mean distance.

### Mean-squared displacement of the EGFR vesicle trajectories

To characterize the motion of EGFR-endosomes in detail, we next analyse the MSDs of trajectory of given *p*_on_, and examine how the on-state velocity *v* and off-state diffusivity *D*_off_ is related to *p*_on_. Because *p*_on_ varies spatially, it is a critical parameter that may reflect the physical and biochemical properties of the vesicle and its interactions with the spatially-varying, local environment. Fig. 4 shows the computed MSDs of the on- and off-state segments from a single cell as a function of delay time *τ*. The MSD curve for the on-state segments is parabolic in the short-time regime and linear in the long-time (Fig. 4(a)). Such a transition from directed motion (short-time) to diffusive motion (long-time) suggests that a randomization of the transport direction in the long-time, which is probably due to that the vesicle performs track-changes and/or constant reversals of transport directions on the same MT track during transport (Fig. S3). We model the on-state transport of EGFR-endosomes to a particle that is propelled with velocity *v* whose direction is subject to an apparent “rotational” diffusion. In this case, the MSD is given as [43]

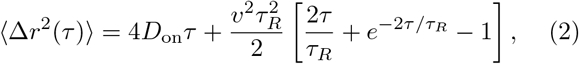

where *D*_on_ is the apparent on-track diffusivity, *τ*_*R*_ is the characteristic coupling time, for which when *τ* ≪ *τ*_*R*_,

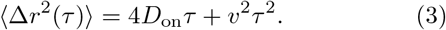

Eq. (2) fits well with the ⟨Δ*r*^2^(*τ*)⟩ for the on-segments with the fitted mean velocity *v* ≃ 0.57 *μm/s* comparable to that obtained from previous studies [7, 24, 39] (see red solid line in Fig. 4(a)), suggesting that this equation catches the essential physics of vesicle’s on-state motion. The off-state motion is not immobile, as compared to the MSDs of immobile QDs that are physically stuck on the coverslip, but diffusive with a diffusion coefficient *D*_off_ ≃ 0.011 *μm*^2^*/s* (Fig. 4(b)). This slow motion may result from the trapping of the EGFR-endosome in a local environment.

**FIG. 4.**
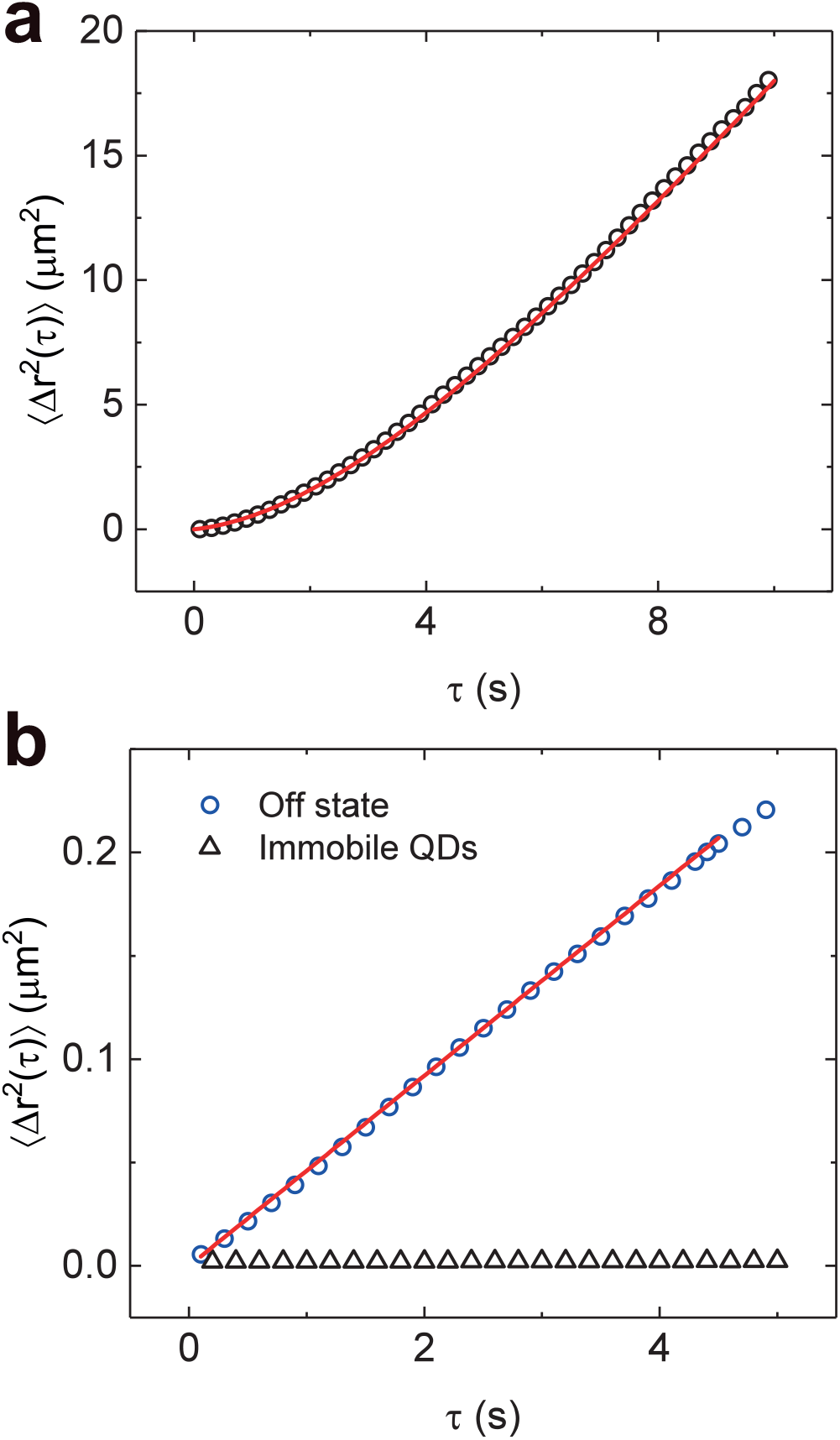
Mean-squared displacement ⟨Δ*r*^2^(*τ*)⟩ of the EGFR vesicle trajectories. (a) Measured ⟨Δ*r*^2^(*τ*)⟩ as a function of delay time *τ* for the on segments of the trajectories from a single cell. The red solid line is a fit of Eq. (2) to the data points with *D*_on_ = 0.06 ± 0.005 *μm*^2^*/s, v* = 0.57 ± 0.02 *μm/s* and *τ* _R_ = 7.3 ± 0.2 *s*. (b) Measured ⟨Δ*r*^2^(*τ*)⟩ for the off segments (blue circles) and immobile QDs (black triangles), which are physically stuck on a coverslip. The red solid line is a linear fit to the data points with *D*_off_ = 0.011 ± 0.002 *μm*^2^*/s*.

Because the fitted *τ*_*R*_ ≃ 7 *s* is much greater than the mean on-state dwell time ≃ 3 *s*, for simplicity, we use Eq. (3) to fit the MSD of all the extracted on-segments from ∼ 80 cells in each bin of *p*_on_ for *τ* ≤ 1 s. The black symbols in Figs. 5 show the fitted *v, D*_on_ and *D*_off_ as a function of *p*_on_. It is clear that the on-track *v* and *D*_on_, and off-track *D*_off_ all increase greatly with *p*_on_. Together with the spatial regulation of *p*_on_ inside the cell (Fig. 3(b)), these results point that as the endosomes approach the perinuclear region, they have a higher probability of being trapped in the off-state, and meanwhile, both their on-state velocity and off-state diffusivity gradually decrease up to 35% and 60%, respectively.

**FIG. 5.**
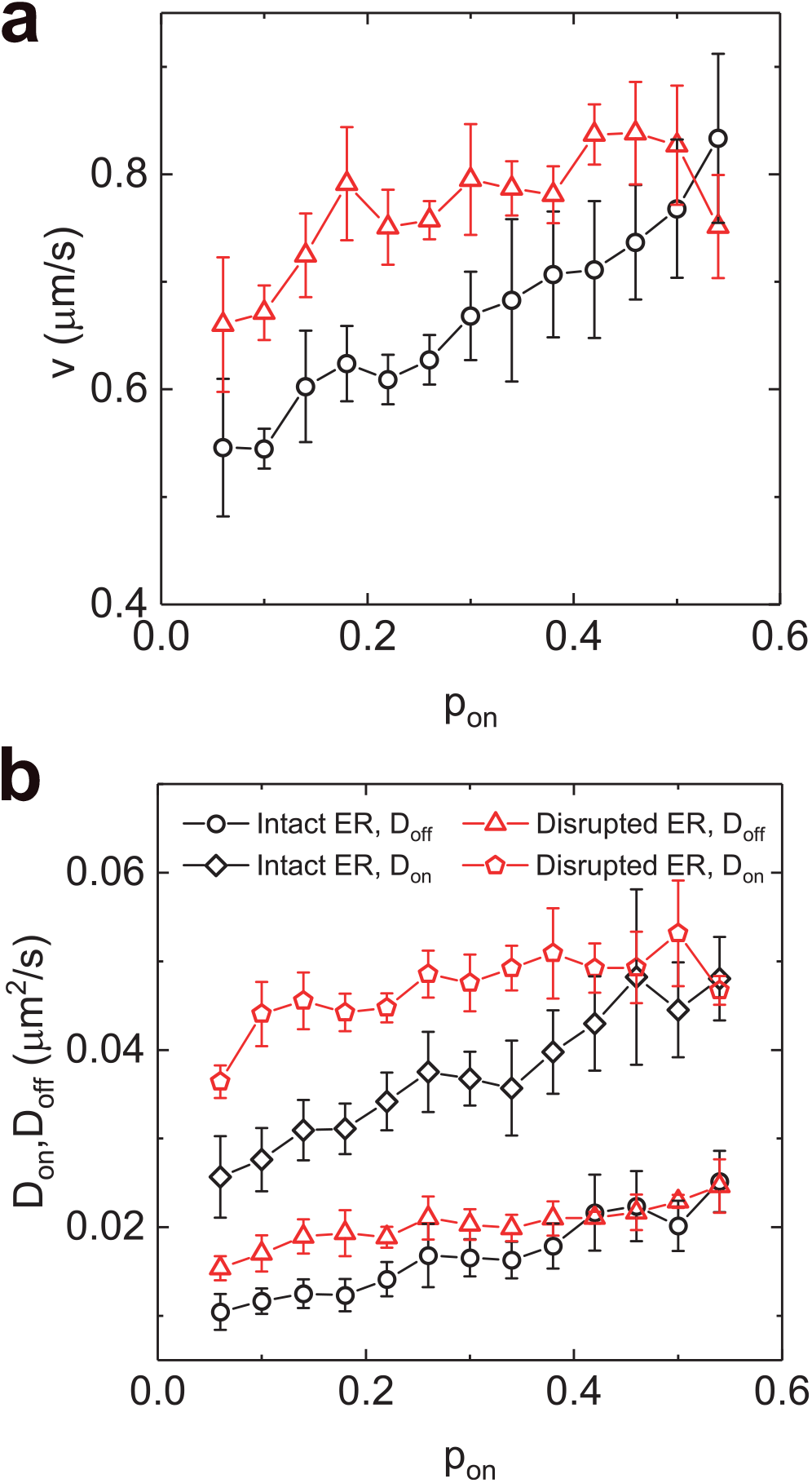
Mean velocity and diffusion coefficients of the EGFR endocytic vesicles. (a) Measured on-state *v* as a function of *p*on for EGFR-endosomes in BEAS-2B cells with intact ER (black symbols) and disrupted ER (red symbols). (b) Measured on-state *D*_on_ and off-state *D*_off_ as a function of *p*on for EGFR-endosomes in BEAS-2B cells with intact ER (black symbols) and disrupted ER (red symbols). *v, D*on and *D*_off_ are obtained by fitting Eq. (3) and ⟨Δ*r*^2^(*τ*)⟩ = 4*D*_off_*τ* to the MSD data points in each bin of *p*on with *τ* ≤ 1 s, respectively. The error bar is the SEM across 4 sets of experiment.

### Dwell time of the on state (*t*_on_) and off state (*t*_off_)

From the extracted large volume of on/off-state segments, we next compute the statistics of the dwell time, *t*_on_ and *t*_off_, for EGFR-endosomes of given *p*_on_ with a bin width = 0.02. The probability density functions (normalized histogram or PDF) of *t*_on_ and *t*_off_ all feature a simple exponential distribution and could collapse onto a single master curve, once the normalized dwell time 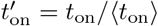 and 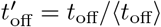are used, where ⟨*t*_on_⟩ and ⟨*t*_off_⟩ are the mean dwell time for endosomes of given *p*_on_ (Figs. 6(a and b)). The red solid lines in Figs. 6(a and b) show the exponential function, which fits the data well, indicating that the transitions between the two states occur randomly in time and could be described by a Poisson process with a constant transition rate [41]. As the vesicle’s *p*_on_ increases, the on-state mean dwell time gradually increases (1.4 ≤ ⟨*t*_on_⟩ ≤ 4.5 s) and the off-state (4.3 ≤ ⟨*t*_off_⟩ ≤ 40.0 s) decreases (Fig. 6(c)). The broader range of ⟨*t*_off_⟩ compared to ⟨*t*_on_⟩ indicates that off-state dynamics are the major contributors of the varied transport behaviours observed for EGFR-endosomes in BEAS-2B cells.

**FIG. 6.**
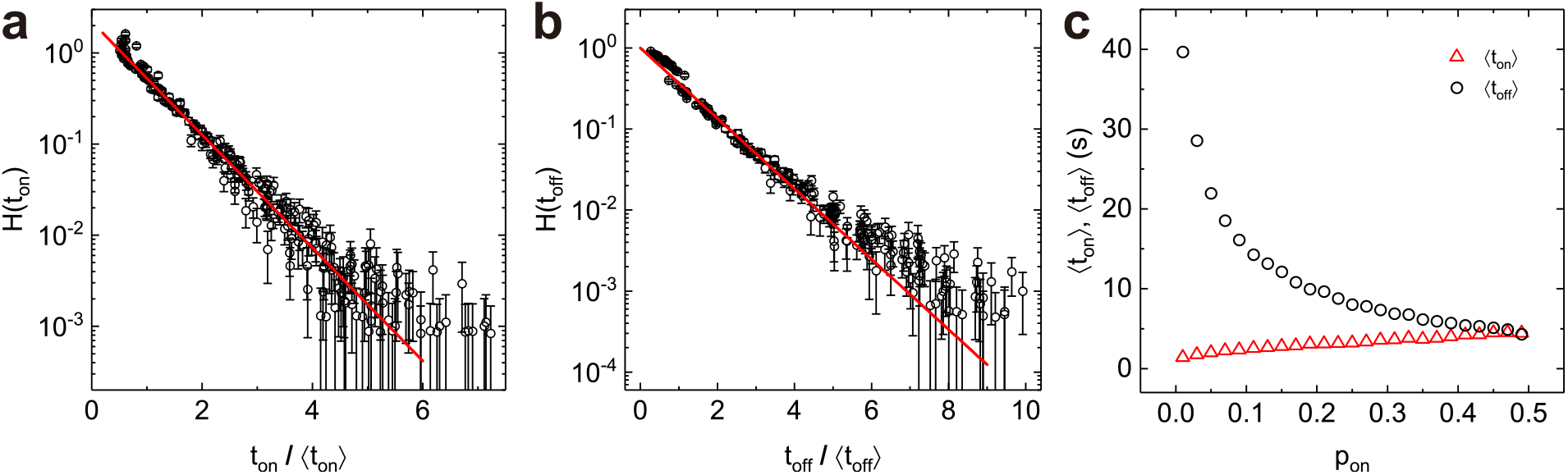
Dwell time of the on state (*t*_on_) and off state (*t*_off_) for EGFR-endosomes of given *p*_on_. (a, b) Measured PDFs *H*(*t*_on_) and *H*(*t*_off_) of the normalized dwell time *t*_on_*/*⟨*t*_on_⟩ and *t*_off_*/*⟨*t*_off_⟩ for trajectories exhibiting different *p*_on_. The red solid lines in (a) and (b) show the exponential functions with *H*(*t*_on_) ≃ 2.2*exp*(−*t*_on_*/*0.7⟨*t*_on_⟩) and *H*(*t*_off_) ≃ *exp*(−*t*_off_*/*⟨*t*_off_⟩), respectively. (c) Measured mean dwell time ⟨*t*_on_⟩ and ⟨*t*_off_⟩ as a function of *p*_on_. A bin width = 0.02 is used to calculate the mean dwell time in each bin of *p*_on_. Overall 10731 trajectories summed from the active fraction of ∼ 80 cells are analysed for the extraction of *t*_on_ and *t*_off_.

In *in-vitro* reconstitution studies, the run-time and run-length of motor proteins on MTs has been found to display a single exponential distribution [44–47], and the pausing time for motors encountering obstacles is also exponentially distributed [48, 49]. Our analysis on the dwell time shows that, in a dynamically heterogeneous media such as a living cell, dwell time for individual EGFR-endosomes both in on and off states are also in the form of a single exponential, *ϕ*_*i*_(*t*) ∝ *exp*(−*t/τ*_*i*_), but subject to a spatially-regulated distribution of mean dwell time *τ*_*i*_. Because MAPs and motor number have been shown to regulate run-length and run-time of motor proteins *in vitro* [17, 47, 50, 51], our observation of the varied on-state mean dwell time (1.4 ≤ ⟨*t*_on_⟩ ≤ 4.5 s) could possibly result from the temporally stable, yet spatially varying distribution of MAPs and/or motor proteins. Qualitative description of pausing behaviours have been previously reported to be spatially correlated with MTs dense region [7], contacts with other endosomes and the ER, our quantification on the off-state mean dwell time which varies up to an order of magnitude indicates that the regulators of the off-state motion also feature a spatial gradient across the cell such as the ER in BEAS-2B cells (Fig. 7(a)). This finding is in agreement with the fact that the ER in unpolarized cells is organized into two distinct structural interconnected domains with a dense flat cisternal sheets occupying the perinuclear region and a less dense reticulated tubules dominating the peripheral regions of cell [28, 31, 32].

**FIG. 7.**
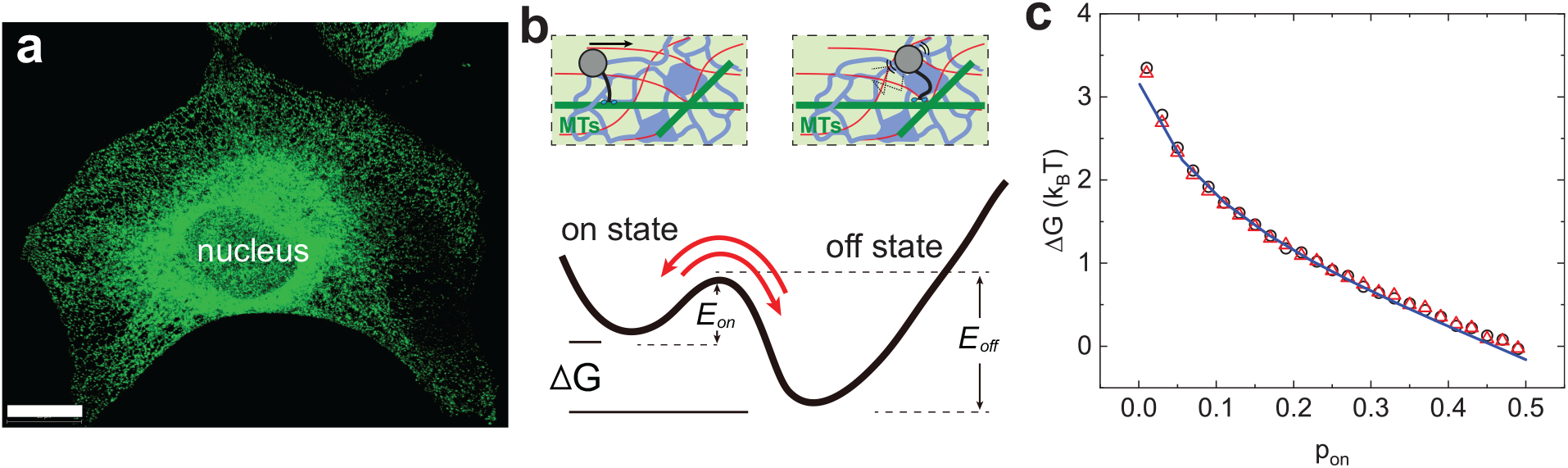
Cargo transport in an energy landscape constructed by ER. (a) 3-dimensional view of the distribution of fluorescently labelled protein disulfide isomerase (PDI, an ER marker) inside a BEAS-2B cell shows that the ER network permeating the whole cytoplasmic space and are highly concentrated at the perinuclear region. Scale bar is 20 *μ*m. (b) Schematic illustration of the 2-state vesicle transport in a double-well potential. The transport in a living cell exists in 2-state: 1) vesicles are carried by motor proteins which move persistently along microtubules (on-state, upper left); 2) directional transport is halted due to the changes of local environment such as microtubule intersections, actin-filament (red) and endoplasmic reticulum (blue) dense region and vesicles perform diffusive motion near the microtubules (off-state, upper right). The transitions between the two states occur randomly in time and one can think of the vesicle is locally in a trapped state with an energy barrier *E*_on_ for the on-state and *E*_off_ for the off-state. (c) Measured Δ*G* as a function of *p*_on_ for EGFR-endosomes in BEAS-2B cells with intact ER (black circles) and disrupted ER (red triangles). The blue solid line shows the fitting with Δ*G* = *Ln*(−(1 − *p*_on_ − *C*)*/*(*p*_on_ + *C*)).

## Discussions

### Theoretical model

To describe the spatially varying on/off dynamics of cargo transport, we propose a simple model that considers vesicles that are carried by motor proteins exist in two states: 1) in the on state, vesicles are attached to the microtubule and motors hydrolyse ATP and perform directional transport with a forward and backward stepping rate, *k*_f_ and *k*_b_; 2) in the off state, vesicles are still attached to the microtubule, however, motors do not hydrolyse ATP and wait for the next ATP hydrolysis with a halt rate, *k*_c_, and vesicles are trapped in the ER network and undergo diffusive motion (Fig. 7(b)). The 2-state model assumes that the transition from on to off occurs continuously and independently at a rate of *k*_f_ + *k*_b_, and hence in a Poisson process. Therefore, the on-state dwell time distribution is simply

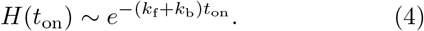

Similarly, the off-state dwell time distribution is

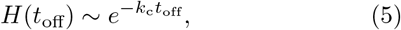

and 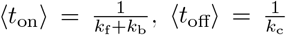. In this simplified two-state system, one can think of the vesicle is locally in a trapped state with an energy barrier *E*_on_ for the on-state and *E*_off_ for the off-state. At steady-state, the transitions are balanced, and we have

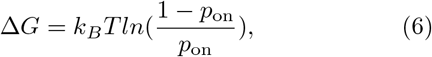

where Δ*G* is the free-energy change. Fig. 7(c) shows the calculated free energy change Δ*G* (black circles), which fits well with Eq. (6) (blue solid line), suggesting that our model catches the essential physics of vesicle’s motion transitions. At the whole cell scale, a large fraction (∼ 80%) of EGFR-endosomes of a small *p*_on_ (*<* 0.15), including non-active endosomes, localize to the perinuclear, ER-rich region, where a high energy trap Δ*G* (*>* 1.5*k*_*B*_*T*) is maintained. For endosomes localizing at the distal region where ER becomes less and less populated, *p*_on_ gradually increases and Δ*G* decreases which favours more on-state behaviours. These results demonstrate the complex EGFR-endosome transport could be described by simple spatially regulated energy landscape Δ*G*(**r**), under which the cell precisely assign the run and pause durations, control the mobility and the steady state distribution of endosomes.

### Role of ER

To validate whether the gradient distribution of ER membrane densities indeed contributes to the spatially varying *p*_on_ and Δ*G*, we first tried to induce ER-stress to disrupt the ER morphology [52]. Acute treatment by immersing the cell in PBS containing BAPTA-AM (1 *μ*M, a cell-permeant chelator of Ca^2+^) for 5 min disrupted the ER morphology drastically. As shown in Fig. 8(a), the ER-marker PDI intensity for the perinuclear ER sheets drops significantly and the tubular ER-network at the cell periphery becomes almost undetectable following the treatment. The altered ER morphology drastically changes the transport dynamics of EGFR-endosomes (Fig. 8(b)), which are evidenced by: (1) the relatively immobile fraction drastically drops to ∼ 19% for cells with disrupted ER (∼ 59% for cells with intact ER, Figs. 8(c), inset), suggesting that ∼ 40% endosomes are mobilized following the ER disruption; (2) as for the mobile fraction, there is a drastic increase (up to ∼ 73%) of high-*p*_on_ endosomes (*p*_on_ *>* 0.15) and drop (up to ∼ 61%) of low-*p*_on_ endosomes, indicating that a large fraction of low-*p*_on_ endosomes changes to high-*p*_on_ endosomes following the treatment (Fig. 8(c)); (3) the measured on-state and off-state mobility *v, D*_on_ and *D*_off_ all show a significant increase with *v* up to ∼ 27%, *D*_on_ up to ∼ 63% and *D*_off_ up to ∼ 57% (Figs. 5(a) and (b), red symbols). Because the normalized dwell time *t*_on_*/*⟨*t*_on_⟩ and *t*_off_*/*⟨*t*_off_⟩ at given *p*_on_ remain exponentially distributed and Δ*G*(*p*_on_) is not sensitive to the treatment (Fig. S4 and Fig. 7(c), red triangles), the altered *H*(*p*_on_) suggests that the energy landscape for endosome transport is shifted to the low-Δ*G* regime which favours more on-state behaviours, and meanwhile, due to the loss of ER and its impedance on the endosome transport, the overall transport is enhanced. Furthermore, the cell becomes more homogeneous following the ER disruption, as evidenced by that the spatial regulation of *p*_on_, and thus Δ*G* especially at the peripheral regions of the cell is lost (Fig. 3(b), red triangles). For endosomes with *p*_on_ *>* 0.2, there is no apparent spatial preference. Similarly, *v, D*_on_ and *D*_off_ become less sensitive to *p*_on_ especially at high-*p*_on_ regime (Figs. 5(a) and (b), red symbols). These results together suggest that an intact ER morphology is required for the maintenance of the observed spatial heterogeneity of endosomes. Increased membrane densities of ER at the perinuclear region not only increase the local crowdedness that may lead to the decreased *D*_off_ for endosomes being in the off state, but also exert increased interactions (or impeding forces) for endosomes being in the on state.

**FIG. 8.**
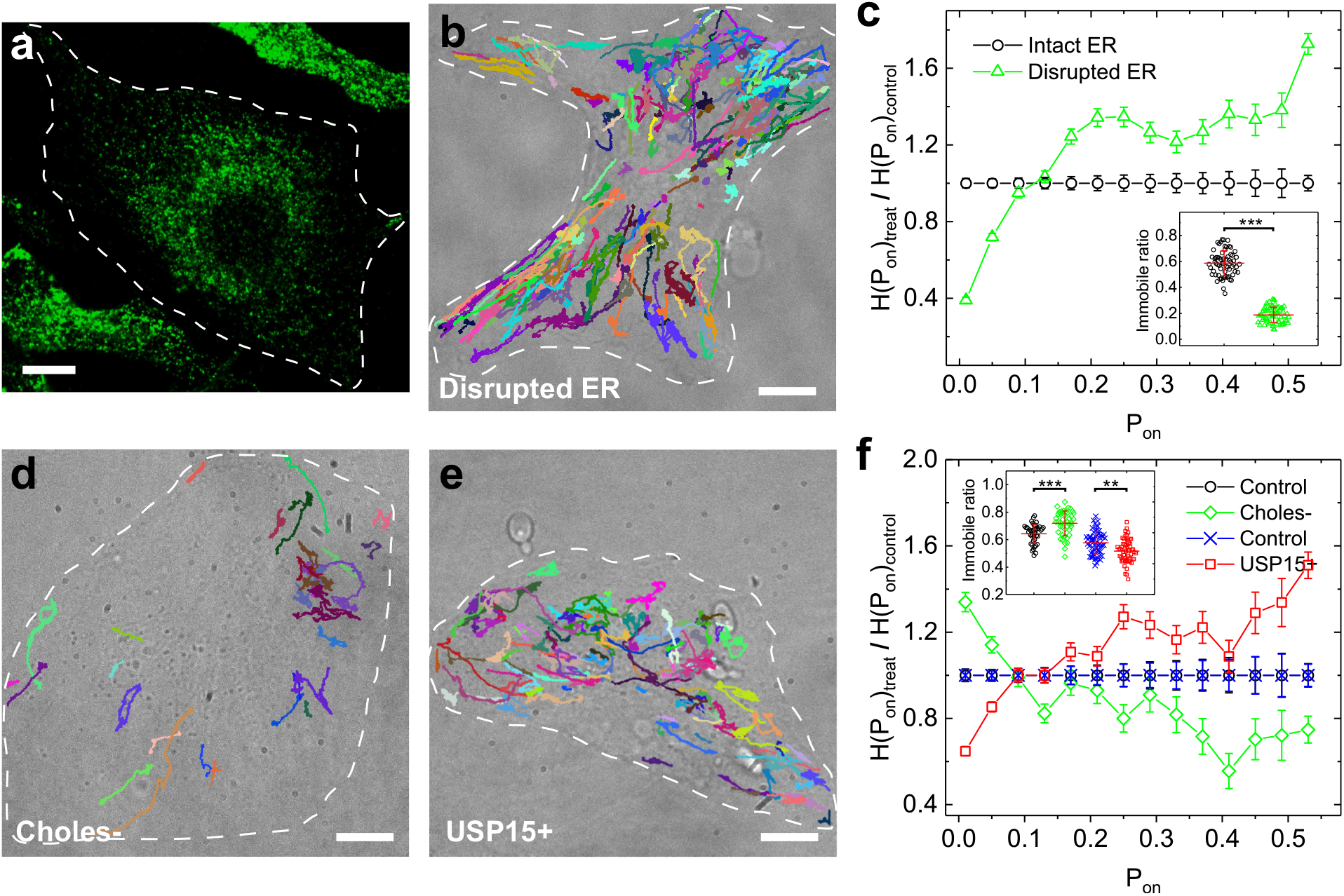
Effects of ER disruption and altered ER-endosome interaction. (a) Fluorescent image of PDI staining shows the ER distribution in BEAS-2B cells after ER-disruption treatment. (b) Overall 175 representative trajectories over the bright-field image with 1800 time steps (180 s) show the dynamics of EGFR-endosomes undergoing active motions in BEAS-2B cells with disrupted ER. (c) Normalized PDF *H*(*p*_on_)*′* for cargo transport in control (intact ER, black circles) and ER-disrupted (green triangles) BEAS-2B cells. Here, *H*(*p*_on_)*′* = *H*(*p*_on_)_*treat*_*/H*(*p*_on_)_*control*_ is used to show the relative ratio-change in comparison with the control groups *H*(*p*_on_)_*control*_. Inset is the comparison of the ratio of the relatively immobile EGFR-endosomes to all the endosomes between cells with intact ER and disrupted ER. *N* = 65 cells for the intact-ER group from 4 independent trials and *N* = 67 cells from the ER-disrupted group from 4 independent trials. ^*\*\*\**^*P <* 0.001. All data points in the inset are plotted with lines indicating means ± S.D.. (d, e) Representative trajectories over bright-field image show the dynamics of EGFR-endosomes undergoing active motions in BEAS-2B cells with either cholesterol-depletion treatment (d, Choles-) or mCherry-USP15 over-expression (e, USP15+). (f) Normalized PDF *H*(*p*_on_)*′* for cargo transport in control, Choles- and USP15+ BEAS-2B cells. Inset is the comparison of the ratio of the relatively immobile EGFR-endosomes to all the endosomes between control and treated cells. *N* = 41 cells for the control group of Choles- and *N* = 44 cells for Choles-group from 3 independent trials, ^*\*\*\**^*P <* 0.001; *N* = 59 cells for the control group of USP15+ and *N* = 54 cells for USP15+ group from 4 independent trials, ^*\*\*\**^*P <* 0.01. Scale bars are 10 *μ*m.

To further test whether ER-specific interactions are involved in the spatial regulation of endosome transport, we next altered the interactions between ER and EGFR-endosomes and evaluated its effects on endosome transport. Two approaches could be conveniently used to alter the ER-endosome interactions according to previous reports: (1) lowering the cholesterol levels in the late endosome membrane could increase the ER-endosome interactions via the late-endosome-localized ORP1L and the ER membrane protein VAP [10, 53]; (2) increasing the cellular level of a deubiquitinating enzyme USP15 could decrease the ER-endosome interactions via acting on the ER-located ubiquitin ligase RNF26 and its associated proteins on endosome membrane [8]. To achieve these purposes, BEAS-2B cells were either pre-depleted of cholesterol (Choles-) by incubating them with methyl-*β*-cyclodextrin (M*β*CD, 2 mM for 30 min) in serum-free buffer solutions [54], before the triggering of the endocytosis of EGFR-endosomes, or overexpressed with red fluorescent protein tagged USP15 (mCherry-USP15, USP15+)[8, 40]. Cholesterol depletion significantly increases the immobile ratio of endosomes, shifts the mobile population towards the low-*p*_on_ regime and decreases the *v, D*_on_ and *D*_off_ during transport (Figs. 8(d), (f) and S5), whereas USP15-mCherry overexpression does the opposite(Figs. 8(e), (f), and S5). Both treatments do not change *H*(*t*_on_), *H*(*t*_off_) and Δ*G*(*p*_on_) (Fig. S6), the altered *H*(*p*_on_) thus suggests that the the energy landscape is shifted to the high-Δ*G* regime and low-Δ*G* regime, respectively, following the modulation of ER-endosome interactions. These results thus demonstrate that ER with its spatially varying morphology and/or with its varied interactions with the endosome could help define a cellular level energy landscape for controlled cargo transport.

The endosomal system in previous studies have been widely described as a bilateral structure with the bulk vesicles quiescently located at the perinuclear region and a small fraction released to the cell periphery [1, 8]. Our current quantitative study, through the extraction of motion states (on and off) and the characterization of the full spectrum of motions at the whole cell scale, presents a physical picture of the system that vesicles in a living cell are travelling in a funnel-shaped energy landscape in which the motion of endosomes are guided towards the perinuclear region (narrow end of the funnel). This funnel-shaped energy landscape is evidenced by that endosomes of given *p*_on_ are found to exhibit a characteristic dwell time either in the on or off state and the spatially varying *p*_on_(**r**) defines the free energy change Δ*G*(**r**) required for endosomes to hop from the off state to the on state.

Further experiments by disrupting the ER and altering the ER-endosome interactions demonstrate that *p*_on_(**r**), and thus Δ*G*(**r**), is largely contributed by the ER network via its spatially varying membrane densities *I*(**r**) (Fig. 7(a)). Increased *I*(**r**) causes increased crowdedness and membrane contacts/interactions between ER and EGFR-endosomes at the perinuclear region and presents a deeper energy trap to hold the endosomes in the off states. Due to the increased ER crowdedness and interactions in this region, *D*_off_ and *v* both decrease gradually (Figs. 5 and S5). Such molecular crowding induced decrease of motion has also been observed in other studies [39, 55]. In this way, cells, via its its whole-cell-spanning ER network, thus could dynamically manipulate the motion states of individual vesicles in space and time to meet various intracellular demands and respond to extracellular signals.

The observed spatial dependence of *p*_on_(**r**) is specific to EGFR-endosomes and likely also specific to Rab7-coated late endosomes or lysosomes that have intense interactions with the ER (data not shown) [2, 8, 10, 30, 53, 56]. For internalized vesicles that does not have specific interactions with the ER such as the Dio-labelled non-specific vesicles, the spatial dependence of *p*_on_(**r**) is lost (Figs. 3(b) and S7). Moreover, vesicles from the immobile fraction are no longer trapped in the perinuclear region but dispersed evenly across the whole cell (Fig. S7).

In the off-state, EGFR-endosomes interact with the ER membrane and perform diffusive motions that are mainly driven by the stochastic, active force fluctuations from the cytoskeleton [12]. How could a off-state endosome escape to the on-state inside the crowded and sticky intracellular environment? We find that this escaping is largely suppressed when the cytoskeletal forces produced by myosin II motors are inhibited by treating the cells with blebbistatin (Bleb) (Fig. S8). Bleb-treatment nearly abolished all high-*p*_on_ endosomes and the immobile ratio increased to ∼ 73% (Fig. S8 (c) and (d)). As myosin-II driven active forces are removed, *D*_off_ drops significantly for ∼ 40-50%, whereas the on-state *v* is less affected (Fig. S8 (e)). These results together suggest that the activity of myosin II motors is essential for the endosome to hop from the off-state to the on-state.

Finally, this study does not exclude the possibility that other cellular factors such as a cellular distribution of the post-translational modifications of tubulin and MAPs may also contribute to the spatial dependence of cargo transport. For example, acetylation of α−tubulin form stable subgroup of MTs and is highly enriched around the MT organizing center which is close to the cell nucleus [57–59]. Whether a localized acetylation could interfere with motor activity and modulate p_on_(r) and ΔG(r) for vesicle transport in a living cell is still largely unknown, which is worthy of future study.

## Supporting information

Supplemental Materials concerning the article: Endosome-ER Interactions Define a Cellular Energy Landscape to Guide Cargo Transport

## Methods

Details about the materials and methods are given in Supplementary Materials.

## Data availability

The data that support the findings of this study (such as figure source data and supplementary information files) are available from the corresponding author (P.T.) upon request.

Acknowledgments

The authors wish to thank Prof. Y. Guo and M. W. for helpful discussions and for providing EGFR antibodies, and R. C. for providing *α*-tubulin antibodies. This work was supported in part by RGC of Hong Kong under grant nos. 16306418 (P.T.), 16302718 (P.T.), 16102417 (P.H.) and AoE/P-02/12 (P.T.) and by MoST of Taiwan under the grant no. 107-2112-M-008-013-MY3 (P.Y.L.).

## Author Contributions

Y.S. and P.T. designed the experiments; Y.S. performed the experiments; Q.Z. constructed the plasmids; all authors analyzed and interpreted data; Y.S. and P.T. drafted the paper with further revisions from all authors; P.T. coordinated the project.

## Competing financial interests

The authors declare no competing financial interests.

## Notes

### Competing Interest Statement

The authors have declared no competing interest.

