## Supplemental Materials concerning the article: Endosome-ER Interactions Define a Cellular Energy Landscape to Guide Cargo Transport for "Endosome-ER Interactions Define a Cellular Energy Landscape to Guide Cargo Transport"

Yusheng Shen[1], Yan Wen[1], Qirui Zhao[2], Pingbo Huang[2], Pik-Yin Lai[3], Penger Tong[1]
*[1] Department of Physics, and [2] Division of Life Science,*
*Hong Kong University of Science and Technology, Clear Water Bay, Kowloon, Hong Kong*
*[2] Department of Physics and Center for Complex Systems,*
*National Central University, Chungli, Taiwan 320, ROC*
(Dated: June 1, 2023)

### I. MATERIALS AND METHODS

**Cell culture, transfection and drug treatment.** BEAS-2B cells, HeLa cells, and HaCaT cells were cultured in Dulbecco's modified Eagle's medium (DMEM, Life Technologies), and NCI-H1975 cells were cultured in RPMI 1640 Medium (Life Technologies). Both media were supplemented with 10% fetal bovine serum (FBS), 50 units/mL of penicillin and 50 µg/mL of streptomycin. All cell cultures were maintained in a 95% air/5% $CO_2$ atmosphere at 37 °C. The cell lines were routinely confirmed to test negative for mycoplasma contamination. For live cell imaging, cells were seeded at a density of approximately $1\times 10^4$ cm$^{-2}$ on a flat or micropatterned glass coverslip which was placed in a 35-mm polystyrene tissue-culture dish.

To inhibit the activity of myosin-II motors, cells were treated with Blebbistatin (10 $\mu$M, Sigma) in culture medium for 15 min before being imaged. To induce ER stress and disrupt ER, cells were first rinsed with phosphate buffered saline (PBS) for three times and then treated with BAPTA-AM (1 $\mu$M, Life Technologies), a cell-permeable chelator for $Ca^{2+}$, in PBS for 5 min before being imaged in PBS for tracking experiments. To increase the intracellular $Ca^{2+}$ level, cells were treated either with ionomycin (0.4 $\mu$M, Sigma), a $Ca^{2+}$ ionophore, or Thapsigargin (1 $\mu$M, Sigma), an inhibitor of the $Ca^{2+}$ ion pump proteins located in the intracellular membranes, in culture medium for 5 min before being imaged in culture medium for tracking experiments. To increase the molecule crowding inside cells, cells were challenged with and imaged in a hypertonic solution (400 mOsm). The control group was treated with an isotonic solution (310 mOsm). These solutions were prepared by adding, respectively, 0.28 and 0.2 M mannitol to a hypotonic base solution (in millimolar: 40 NaCl, 5 KCl, 1 $CaCl_2$, 2 $MgCl_2$, 10 HEPES (pH 7.4), 91 mOsm) to maintain a constant ionic strength. To increase ER-endosome interactions, cells were treated with methyl-$\beta$-cyclodextrin (M$\beta$CD, 2 mM for 30 min) in serum-free, isotonic buffer solutions. Cells incubated in serum-free, isotonic buffer solutions without adding M$\beta$CD are used as control. All the treatments were performed 15 min after the labelling of EGFR with EGF-QDs.

The expression vectors used in the study were pmCherry-C1 and mCherry-USP15. For generating mammalian expression mCherry-USP15, human USP15 complementary DNAs (Flag-HA-USP15, 22570, addgene) were amplified and subcloned into the pmCherry-C1 expression vector. Transfections were performed by using the Lipofectamine 2000 reagent kit (Invitrogen) according to manufacturer's instructions.

**EGFR labelling, Dio labelling and optical imaging.** The stock EGF-QD complex was prepared by mixing biotin-EGF (4 nM, Thermo Fisher Scientific) with streptavidin-conjungated quantum dots (QDs) with maximum fluorescent emission at 655 nm (QD655, 2 nM, Thermo Fisher Scientific) in vitro in culture medium at 4 °C for 3 h. For high speed, short duration recording of EGFR-endosomes, EGFRs in the plasma membrane were labelled with EGF-QDs at a concentration of 0.8 nM biotin-EGF at room temperature for 5 min. Nonspecific vesicles are labelled by incubating BEAS-2B cells in culture medium containing the lipophilic membrane dye $DiOC_{18}(3)$ (DiO, 5 $\mu$M, Invitrogen) at 37 °C for 30 min to allow nonspecifically labelled cell membranes to be internalized into the cell. After washing with culture medium for three times (5 min each), the cell-containing glass coverslip was then mounted on a coverslip holder (SC15012, Aireka Cells), which was finally mounted on an inverted microscope (DM-IRB, Leica) with a 100× objective (NA = 1.40). A live cell imaging chamber (CU-501, Chamlide TC) was equipped to the microscope to provide optimal culture conditions (95% air/5% $CO_2$ atmosphere at 37 °C) for cells during imaging.

Image sequences were typically recorded at 10 fps (frame per second) for 5 min or 1.33 fps for 1 h by using an electron-multiplying charged-coupled device (EMCCD) camera (Ixon3 897, Andor). The exposure time for each frame is 30 ms and a microscope shutter was used to control the overall UV exposure time and reduce the light-induced damage to the living cells. The recorded images have 16 bits of gray scales and a spatial resolution of 512×512 pixels with the width of each pixel $p_w = 133$ nm in our optical setup. As described previously and also partially shown in the inset of Fig. 3a, the accuracy of the measurement of the QDs displacements was determined from the recording of stationary QDs that were stuck on the glass coverslip. Their total displacement over a time period of 300 s is less than 24 nm, which is much less than the typical displacement of EGFR-endosomes.

**Immunostaining and confocal microscopy.** The entire staining process was performed at room temperature. Briefly, BEAS-2B cells were fixed with 4% paraformaldehyde in PBS (for 10 min) and subsequently permeabilized with 0.2 M $NH_4Cl$/PBS containing 0.2% Triton X-100 (for 10 min), and then blocked with 4% bovine serum albumin (Sigma-Aldrich) in PBS (for 2 h). To visualize the spatial distribution of PDI, $\alpha-$tubulin and internalized EGFRs, the cells were first incubated with mouse monoclonal antibody against PDI (1:200, ADI-SPA-891, Enzo Life Sciences), or $\alpha-$tubulin (1:200, T6199, Sigma-Aldrich), or EGFR (1:50, E2906, Sigma-Aldrich) for 1 h. After washing with PBS (5 min each for three times), the cells were incubated with Alexa Fluor 488 conjugated goat anti-mouse IgG (1:200 dilution, Abcam) for 1 h. Finally, the cell sample was washed with PBS (5 min each for three times) and mounted onto a microscope slide with Citifluor-AF-1 (Ted Pella), and then examined under a confocal microscope (TCS SP8, Leica). The fluorescent images were analyzed by using ImageJ software (NIH).

**Single-particle tracking and extraction of on/off-state segments.** Single-particle tracking (SPT) was performed using a homemade tracking program written in MATLAB as previously described [1], which is based on the standard tracking algorithm [2, 3]. With this advanced SPT algorithm, we were able to obtain the position $\mathbf{r}(t)$ at time $t$ for EGFR-endosomes and Dio-labeled non-specific vesicles, and their trajectories were constructed from the consecutive images.

To extract the on/off (directed/diffusive) state segments from the individual trajectories, an automatic algorithm that could characterize the degree of local directionality of movements was used [4] (Fig. S2). We analyzed the turning angles, $\theta$, around an arbitrary point in a given trajectory at different time scales (5 time steps in this study) and discretize them into 1 if $\theta < \pi/2$ (forward motion) and 0 if $\theta \geq \pi/2$ (backward motion). After averaging the discretized angles, a threshold at 0.5 is used to segment the trajectory into on ($\langle\theta\rangle \geq 0.5$) and off states ($\langle\theta\rangle < 0.5$). Finally, a sliding window (10 time steps, 1 s) was applied to filter out the transient hopping events between on and off states to resolve the final on and off segments. Only on-state events longer than 0.3 s were considered as true on-state.

To obtain the off-state diffusion coefficient $D_{\text{off}}$, mean squared displacements (MSDs), $\langle\Delta\mathbf{r}^2(\tau)\rangle = \langle(\mathbf{r}(t+\tau) - \mathbf{r}(t))^2\rangle$, were computed from the off-state segments and fitted to $\langle\Delta\mathbf{r}^2(\tau)\rangle = 4D\tau$ to obtain the diffusion coefficients of EGFR-endosomes.

### II. SUPPLEMENTARY FIGURES AND TABLES


[1] Shen, Y. *et al.* Directed motion of membrane proteins under an entropy-driven potential field generated by anchored proteins. *Phys. Rev. Res.* **3**, 043195 (2021).

[2] Crocker, J. C. & Grier, D. G. Dethods of digital video microscopy for colloidal studies. *J. Colloid Interface Sci.* **179**, 298-310 (1996).

[3] Anthony, S., Zhang, L. & Granick, S. Methods to track single-molecule trajectories. *Langmuir* **22**, 5266-5272 (2006).

[4] Röding, M., Guo, M., Weitz, D. A., Rudemo, M. & Särkkä, A. Identifying directional persistence in intracellular particle motion using Hidden Markov Models. *Math. Biosci.* **248**, 140-145 (2014).

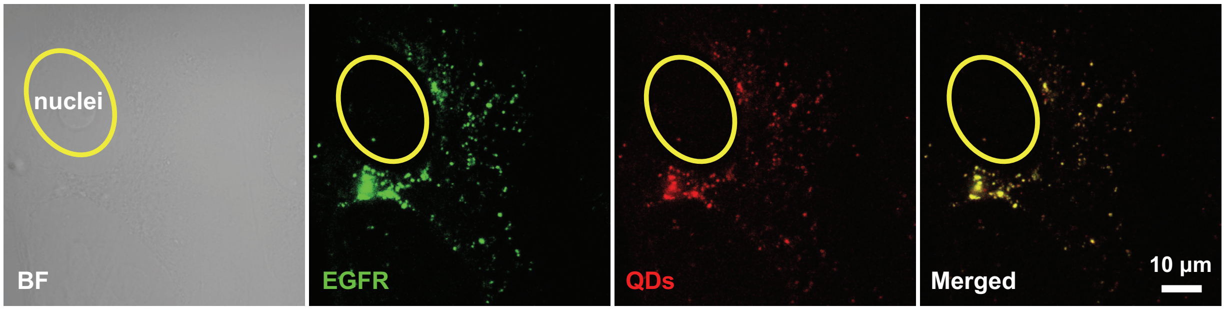


FIG. S1. Internalized EGFRs colocalize with QDs signal. Fluorescent images of BEAS-2B cells stained for EGFR (green) after the endocytosis of EGF-Qds (red). Yellow circle indicates the position of nucleus. Scale bar is 10 $\mu$m.

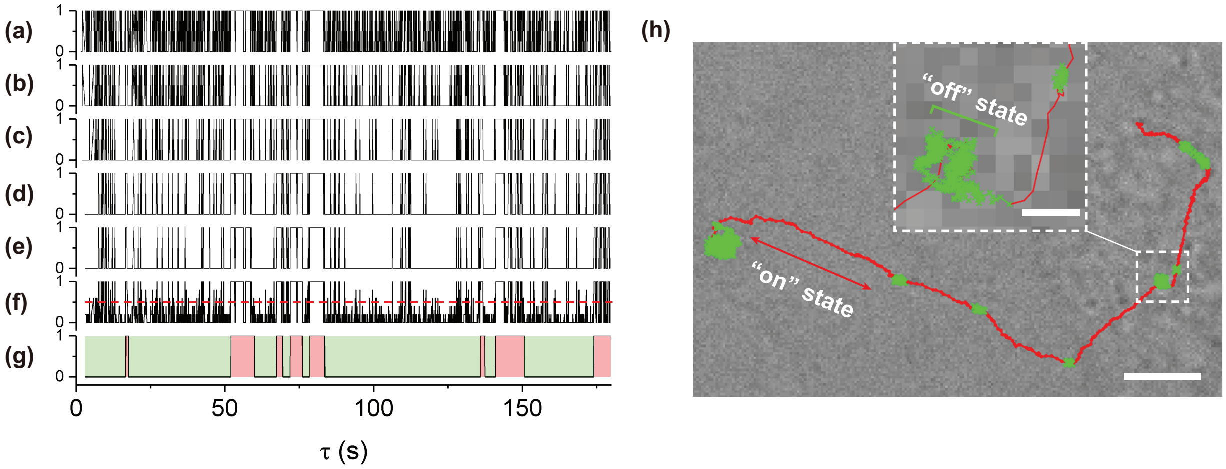


FIG. S2. **Analysing method on the on/off dynamics of cargo transport.** (a-e) Discretized turning angles around an arbitrary point in the trajectory, as shown in (h), at different time scales (from top to bottom are 0.1 $s$, 0.2 $s$, 0.3 $s$, 0.4 $s$ and 0.5 $s$). The calculated turning angles, $\theta$, ranging from 0 (straight forward motion) to $\pi$ (straight backward motion), regardless of the left and right turns, are assigned as 1 if $\theta < \pi/2$ and 0 if $\theta \geq \pi/2$. (f) Averaged discretized angles from (a-e). A threshold at 0.5 (red dashed line) is used to segment the trajectory into on ($\geq$ 0.5) and off states ($<$ 0.5). (g) Distinct on (red) and off (green) states are finally extracted by using a sliding window to filter out the transient hopping events between on and off states. (h) A representative trajectory shows the switching between the on state (red) and the off state (green). Inset shows a magnified view of a portion of the trajectory. Scale bars are 5 $\mu$m, and 0.5 $\mu$m for the inset.

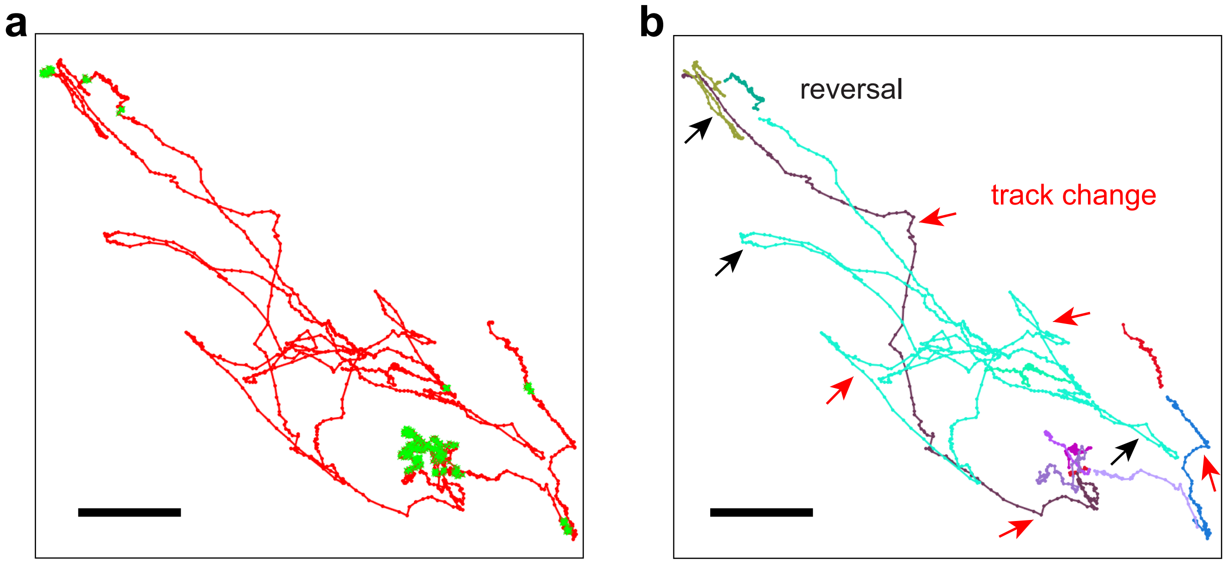


FIG. S3. (a) An example trajectory shows the transitions between on (red) and off (green) states for EGFR endosomes. (b) Extracted on-state segments (coloured trajectories) of the trajectory shows the endosome performs track-changes (indicated by red arrows) and constant reversals of transport directions on the same MT track (indicated by black arrows). Scale bars are 2 $\mu$m.

| Cell type | Cell treatment | No. of cells | No. of active trajectories |
|---|---|---|---|
| BEAS-2B | EGFR labelling | 83 | 10731 |
| BEAS-2B | EGFR labelling; ER disruption | 66 | 10132 |
| BEAS-2B | Cholesterol depletion control; EGFR labelling | 41 | 7068 |
| BEAS-2B | Cholesterol depletion; EGFR labelling | 44 | 3978 |
| BEAS-2B | mCherry transfected control; EGFR labellingl | 59 | 7657 |
| BEAS-2B | mCherry-USP15 transfected; EGFR labelling | 54 | 8081 |
| BEAS-2B | DiO labelling | 54 | 17347 |
| BEAS-2B | EGFR labelling; Blebbistatin treatment | 68 | 5406 |
| | Total: | 469 | 70400 |

TABLE I. Summary of cell treatments and sample sizes. The columns, from left to right, list the type of cells used, the treatments to the cells, the number of cells imaged, and the total number of active trajectories used in the statistical analysis, respectively.

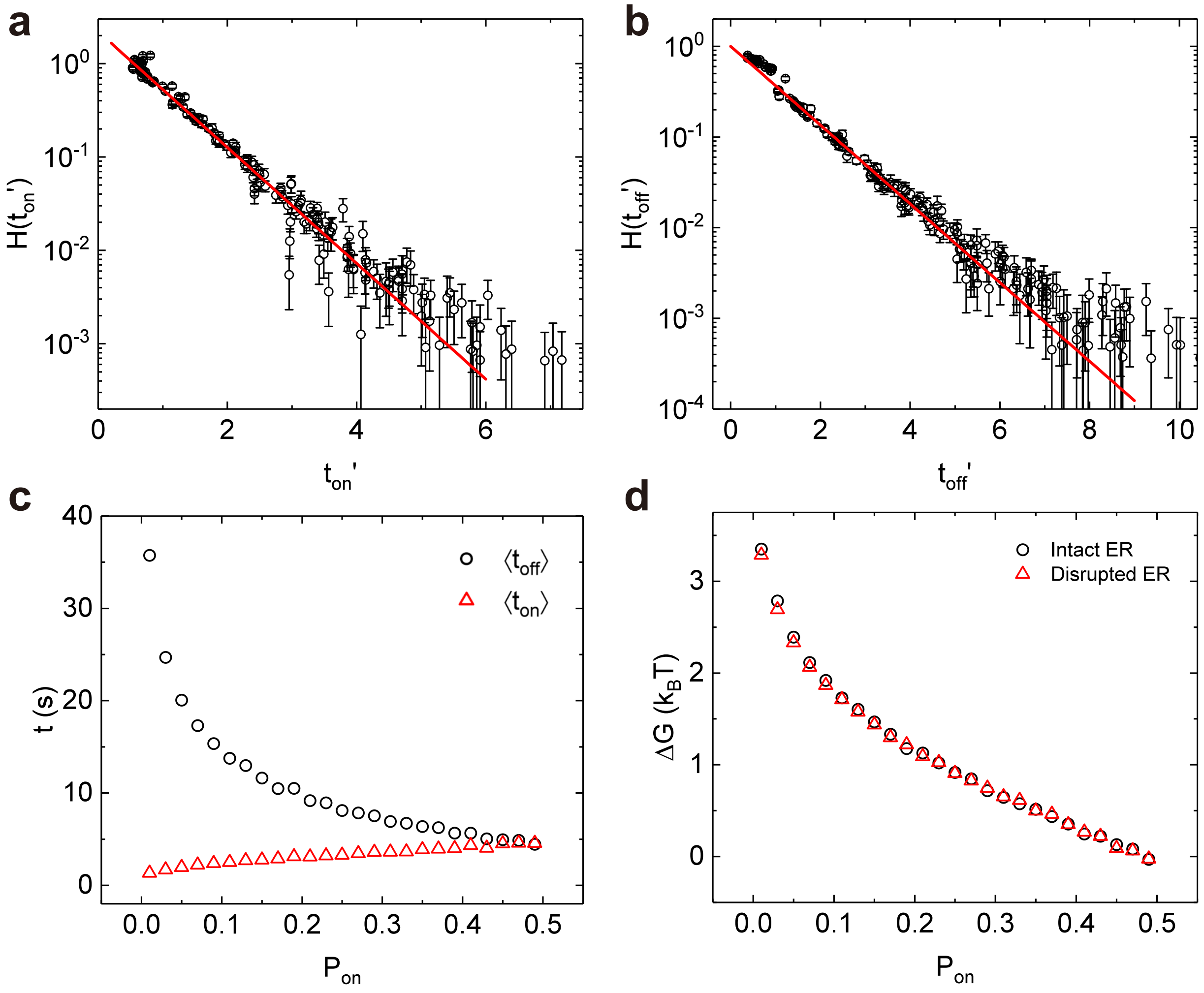


FIG. S4. **ER disruption does not alter the statistical property of $t_{\rm on}$ and $t_{\rm off}$, and $\Delta G(p_{\rm on})$.** (a, b) Measured PDFs $H(t'_{\rm on})$ and $H(t'_{\rm off})$ of the normalized dwell time $t'_{\rm on}$ and $t'_{\rm off}$ for trajectories exhibiting different $p_{\rm on}$. The red solid lines in (a) and (b) show the exponential functions with $H(t'_{\rm on}) \simeq 2.2exp(-t'_{\rm on}/0.7)$ and $H(t'_{\rm off}) \simeq exp(-t'_{\rm off})$, respectively. Overall 10132 trajectories summed from the active fraction of $\sim$ 66 cells are analysed for the extraction of $t_{\rm on}$ and $t_{\rm off}$. (c) Measured mean dwell time $\langle t_{\rm on}\rangle$ and $\langle t_{\rm off}\rangle$ as a function of $p_{\rm on}$. (d) Measured $\Delta G$ as a function of $p_{\rm on}$ for BEAS-2B cells with intact ER (black circles, also shown in Fig. 7c) or with disrupted ER (red triangles).

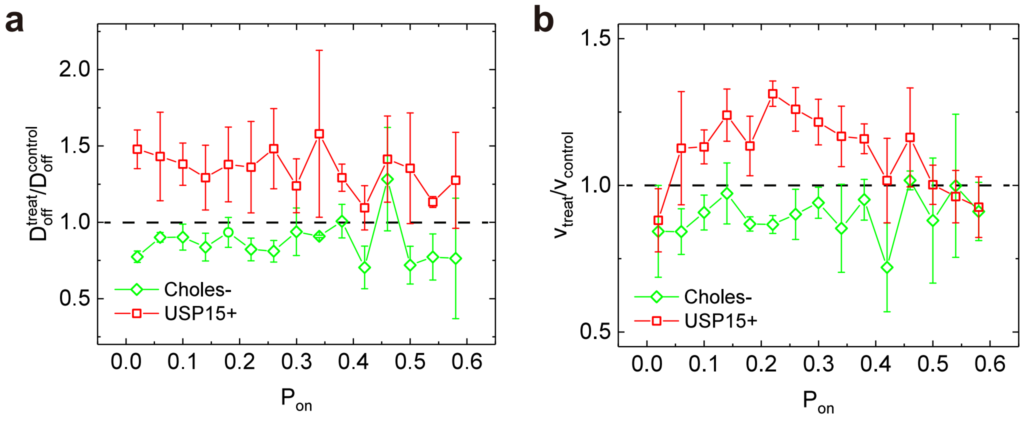


FIG. S5. **Effects of altered ER-endosome interaction on $D_{\rm off}$ and $v$.** Relative change of off state diffusion coefficient $D^{treat}_{\rm off}/D^{control}_{\rm off}$ (a) and on state velocity $v_{treat}/v_{control}$ (b) as a function of $p_{\rm on}$. The error bar is the SEM across 3 sets of independent experiment. Black dashed lines are added to guide the eye for comparison.

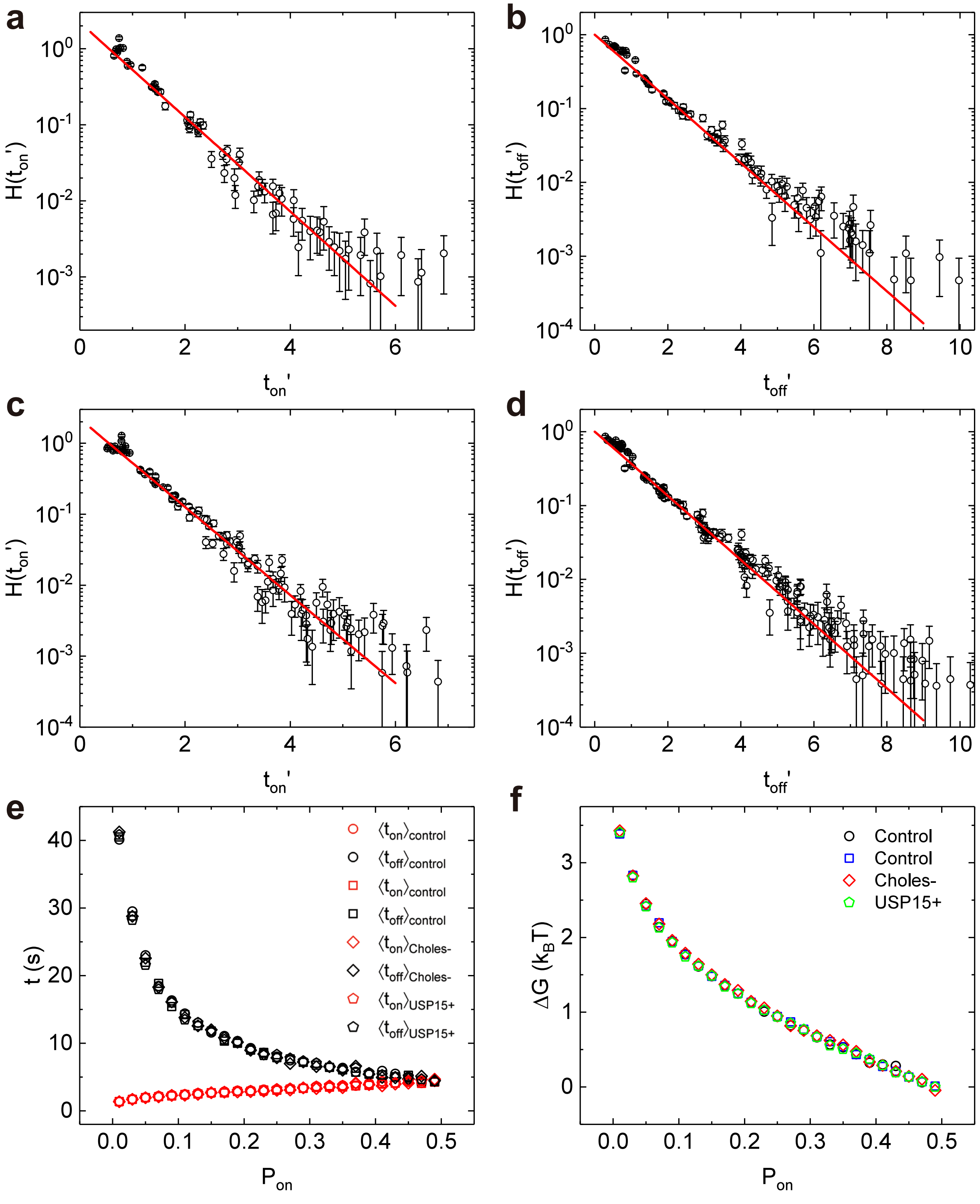


FIG. S6. **Altered ER-endosome interactions do not change the statistical property of $t_{\rm on}$ and $t_{\rm off}$, and $\Delta G(p_{\rm on})$.** (a-d) Measured PDFs $H(t'_{\rm on})$ and $H(t'_{\rm off})$ of the normalized dwell time $t'_{\rm on}$ and $t'_{\rm on}$ for trajectories exhibiting different $p_{\rm on}$. For simplicity, only the data from cholesterol-reduced (a, b) and mCherry-USP15 expressed (c, d) cells are plotted here for demonstration. The normalized dwell time $t'_{\rm on}$ and $t'_{\rm off}$ in control cells also display exponential distributions. The red solid lines in (a, c) and (b, d) show the exponential functions with $H(t'_{\rm on}) \simeq 2.2exp(-t'_{\rm on}/0.7)$ and $H(t'_{\rm off}) \simeq exp(-t'_{\rm off})$, respectively. Details about the sample sizes are summarized in Table S1. (e) Measured mean dwell time $\langle t_{\rm on}\rangle$ and $\langle t_{\rm off}\rangle$ as a function of $p_{\rm on}$ for BEAS-2B cells under different treatments.. (f) Measured $\Delta G$ as a function of $p_{\rm on}$ for BEAS-2B cells under different treatments.

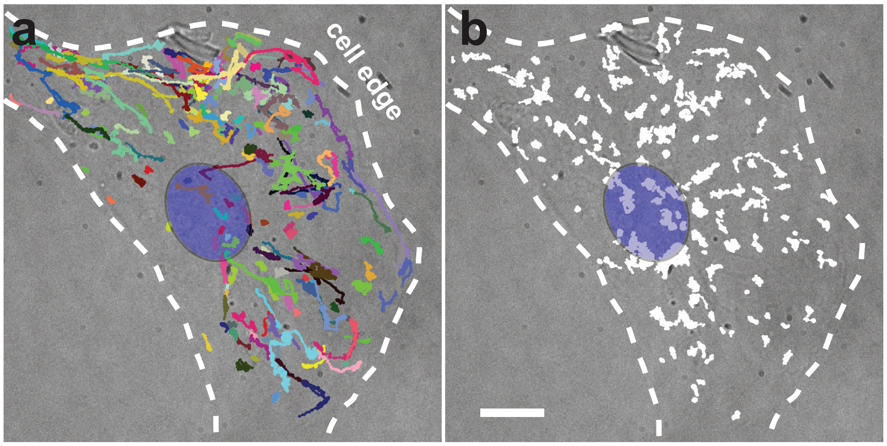


FIG. S7. **Dio-labelled nonspecific vesicles display no spatial dependence of $p_{\mathrm{on}}$.** Overall 465 representative trajectories with 1800 time steps (180 s) show the dynamics of Dio-labelled nonspecific vesicles inside a living BEAS-2B cell. The active (coloured, $N = 190$ trajectories) and non-active (white, $N = 275$ trajectories) trajectories over bright-field images are shown in (a) and (b), respectively. The blue ellipse indicates the nucleus and the white dashed lines indicate the cell edge. Scale bar is 10 $\mu$m.

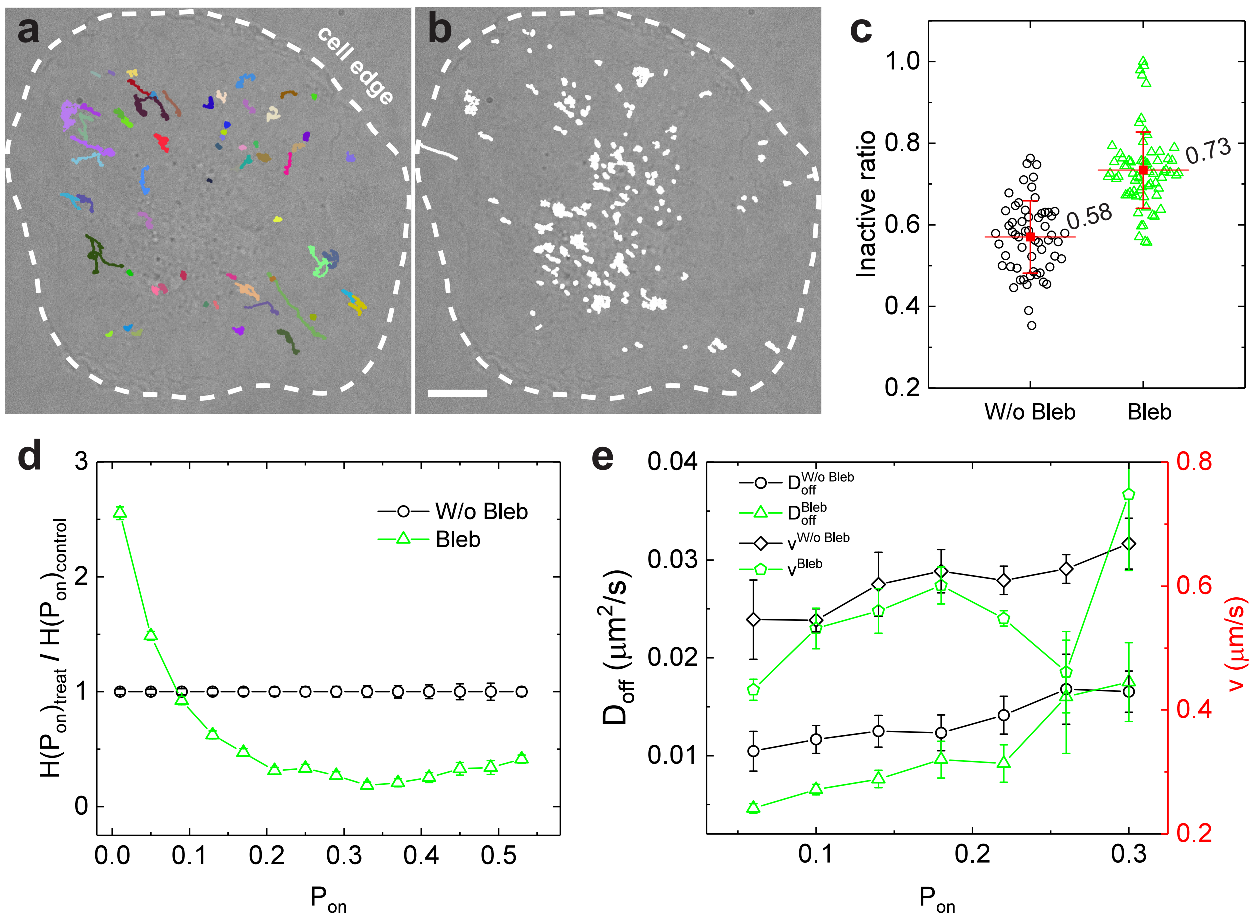


FIG. S8. **Active force fluctuations from the cytoskeleton are required to drive the hopping of EGFR-endosomes from off-state to on-state.** (a, b) Overall 284 representative trajectories with 1800 time steps (180 s) show the dynamics of EGFR-endosome inside a living BEAS-2B cell following Blebbistatin treatment (Bleb). The active (coloured, $N = 63$ trajectories) and non-active (white, $N = 221$ trajectories) trajectories over bright-field images are shown in (a) and (b), respectively. The white dashed lines indicate the cell edge. Scale bar is 10 $\mu$m. (c) Comparison of the inactive ratio of EGFR-endosomes between cells with Bleb treatment (green triangles) and without (W/o) Bleb treatment (black circles). (d) Normalized PDF $H(p_{\rm on})'$ as a function of $p_{\rm on}$ for EGFR-endosome transport in BEAS-2B cells with or without Bleb treatment. (e) Comparison of measured $D_{\rm off}$ and $v$ in BEAS-2B cells with (green symbols) or without (black symbols) Bleb treatment. Because Bleb treatment nearly abolished endosomes for $p_{\rm on} > 0.2$, only low-$p_{\rm on}$ endosomes are compared here. The data for cells without Bleb treatment is reproduced from Fig. 3b. The error bar is the SEM across 3 sets of experiment.
